## Supplementary Figures 1-11 and Tables 1-4 for "Ion currents through Kir potassium channels are gated by anionic lipids"

Jin *et al.*

### Figures

Supplementary Figure 1. Kir Sequence alignment and ACMA assay.
Supplementary Figure 2. Crystal structures show the L124M mutant retains structural integrity.
Supplementary Figure 3. Single channel recordings of L124M show a preference for the fully open state.
Supplementary Figure 4. Comparative PMFs at the Leu124 and Tyr132 collars
Supplementary Figure 5. Ions only pass the Leu124 collar when the aperture has sufficiently widened.
Supplementary Figure 6. The *gauche* rotamer of Tyr57 blocks lipid entering the fenestrations
Supplementary Figure 7. MTS-alkyl derivatization.
Supplementary Figure 8. Normalised data from liposomal fluorescence flux (ACMA) assays.
Supplementary Figure 9. HMM analysis of single channel recordings of C119-alkylated KirBac3.1
Supplementary Figure 10. Aliphatic chain occupancy affects pore aperture in C119-derivatised channels.
Supplementary Figure 11. Pairwise cross comparison of the probability density distribution of PC, PS and PG in Fig. 4.

### Tables

Supplementary Table 1. Crystallographic Structure Determination and Refinement.
Supplementary Table 2. Summary of Molecular Dynamics Simulations.
Supplementary Table 3. Summary statistical data on fluorescence change for functional ACMA assays
Supplementary Table 4. Summary rates data for ACMA assays.

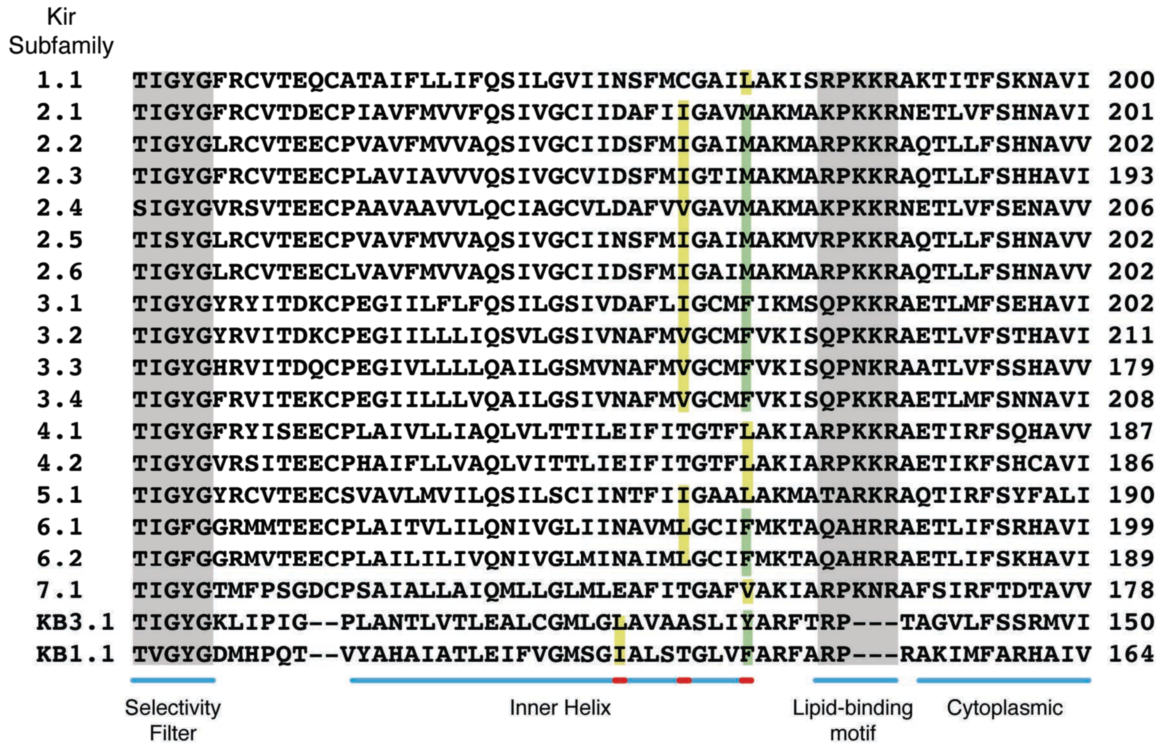


**Supplementary Figure 1**. **Kir Sequence alignment and ACMA assay.** Partial sequence alignment of human Kir subfamily members against the prokaryotic homologues KirBac1.1 (*Burkholderia pseudomallei*) and KirBac3.1 (*Magnetospirrilum magnetotacticum*). Labelled blue underscoring shows the sequence-to-structural alignment, with red segments indicating the inward-facing residues, each a helix-turn apart, lining the conduction pathway. The selectivity filter and basic lipid binding motif are shaded grey. Green shading indicates the residue constricting the cytoplasmic opening at the helix bundle crossing (the conventional site of the 'gate') and yellow the inward facing branched aliphatic residues – all have only one potential constriction site, except Kir5.1, which has two.

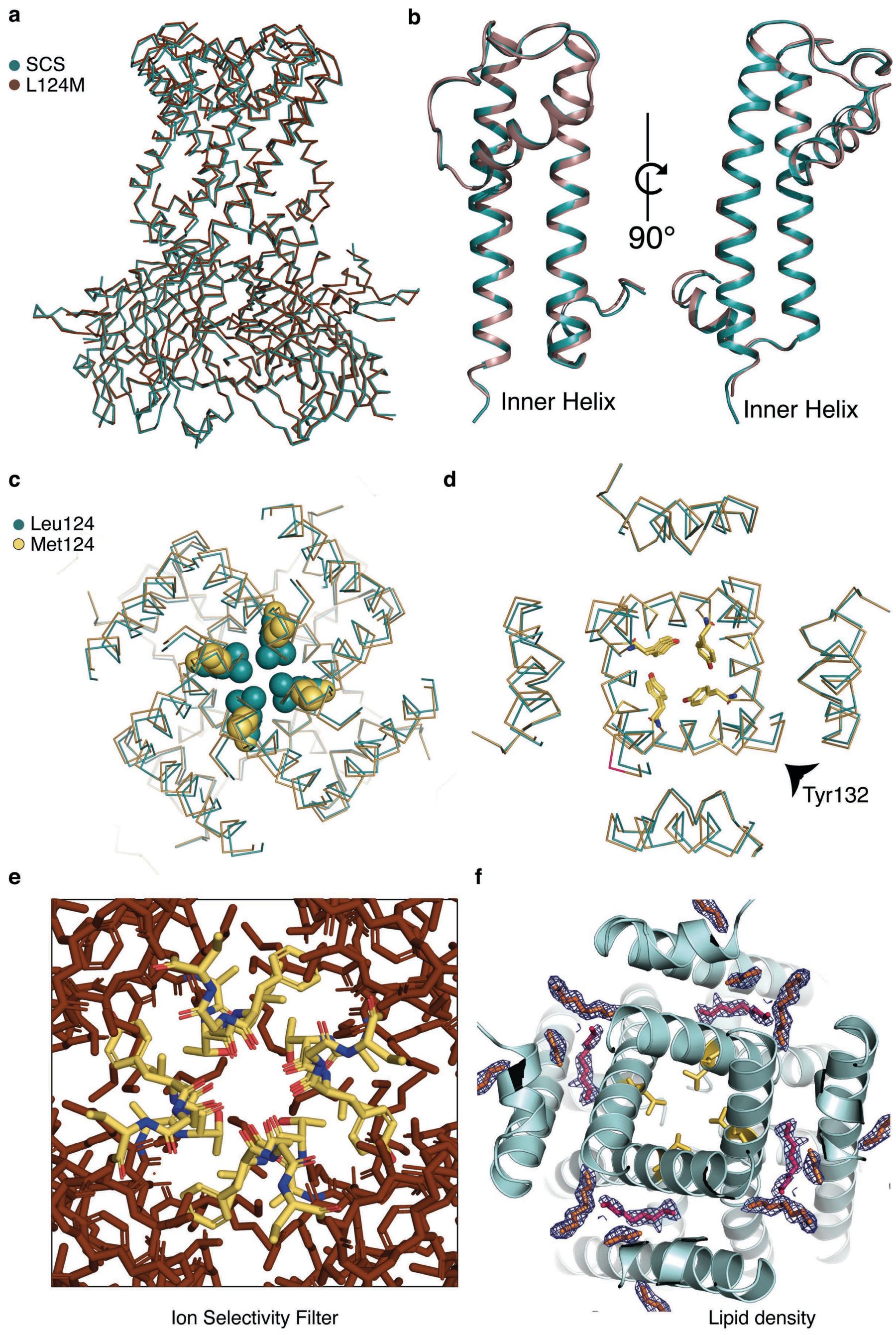


**Supplementary Figure 2**. **Crystal structures show the L124M mutant retains structural integrity**. (**a**) Ca overlay of the entire polypeptide backbones of the KirBac3.1-SCS (cyan) and L124M (SCS-L124M; brown). (**b**) Orthogonal views of a single subunit of KirBac3.1-SCS superposed on L124M. (**c**) cross-sectional slice of the overlay at Leu124 shows that, despite different side chains at residue 124, the Ca backbone does not significantly widen. (**d**) Cross-sectional slab of the overlay at the helix bundle constriction Tyr132. (**e**) The selectivity filter (TVGYG) in the L124M structure is conserved, with all selectivity filter carbonyls pointing inwards and coordinating ions (not shown). (**f**) Several lipid fragments are built into the structure; a 2.4 Å sharpened (B factor -50 $Å^2$) ($2|F_o|-|F_c|$ electron density map is contoured at 1 $\sigma$ (dark blue mesh).

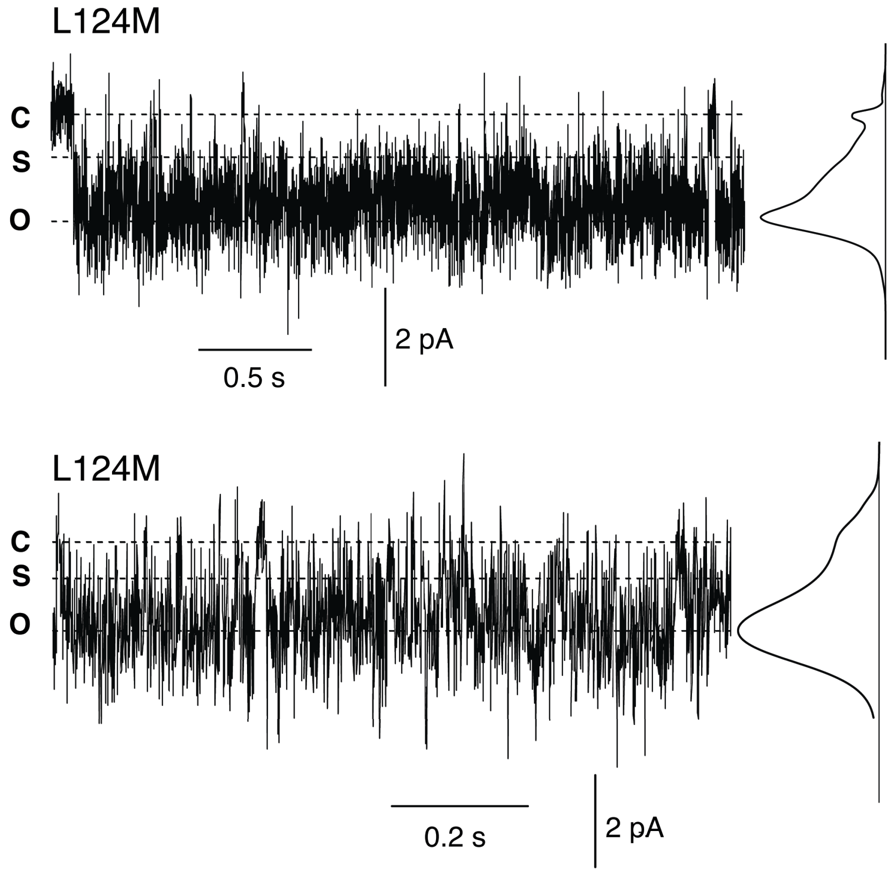


**Supplementary Figure 3. Single channel recordings of L124M show a preference for the fully open state.** Single channel recordings of the L124M mutant of KirBac3.1 at -60 mV (top) and -70 mV (bottom), filtered at 500 Hz. Open (O), closed (C) and partially open substate (S) levels determined by Hidden Markov Models are indicated by horizonal dashed lines. The maximum-likelihood amplitude histograms generated by Hidden Markov Models are aligned with the recordings.

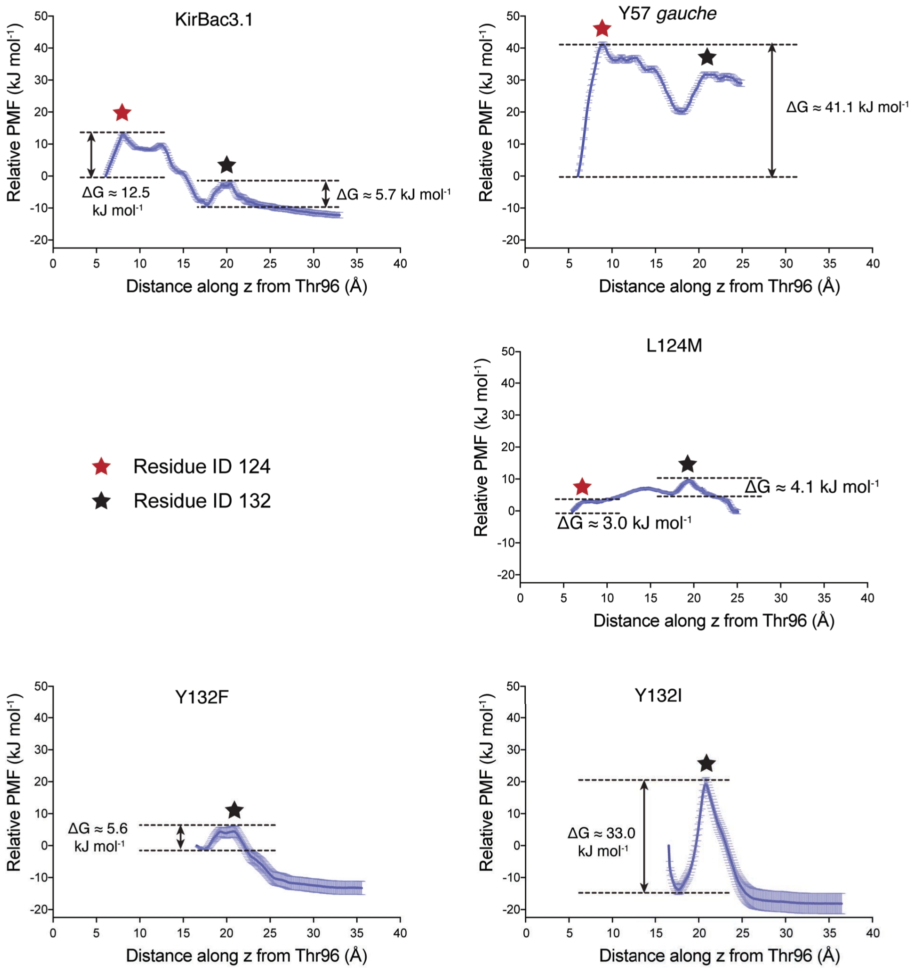


**Supplementary Figure 4. Comparative PMFs at the Leu124 and Tyr132 collars.** This shows the free energy barrier faced by a permeating $K^+$ at Leu124 collar and/or Tyr132 collar, in point mutants and in channels with the fenestration-blocking rotamer of Tyr57. The potential of mean force along the molecular 4-fold axis of a wild type KirBac3.1 structure is shown as a function of the distance between a $K^+$ cavity ion 'pulled' along the *z*-axis and the centre of mass of the four Thr96 sidechains (the global reaction coordinate (see Figure 1a)). SEM are depicted as vertical lilac bars.

**a**

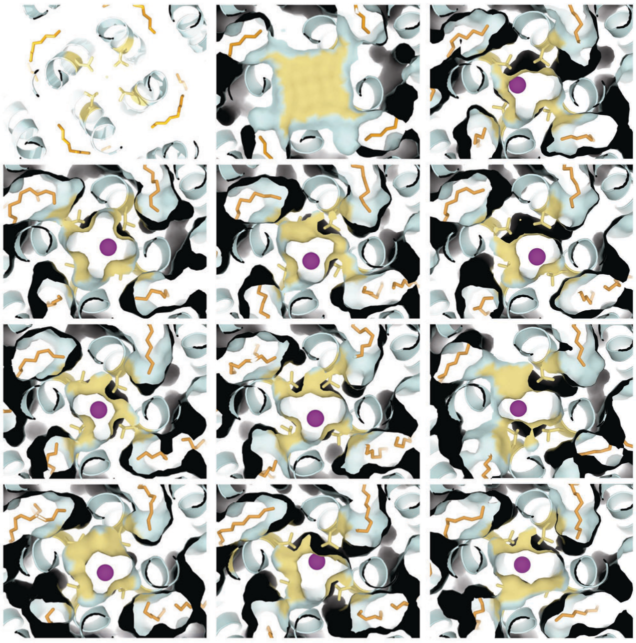

**b**

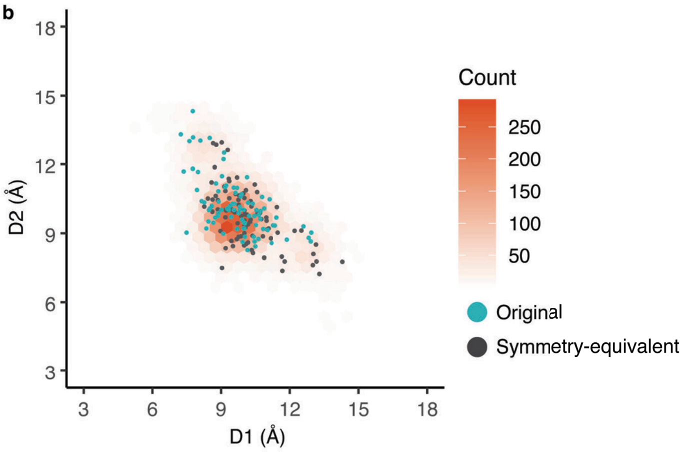


**Supplementary Figure 5. Ions only pass the Leu124 collar when the aperture has sufficiently widened**. (**a**) Cross-sectional slice through the KirBac3.1 pore (cyan) at the Leu124 collar of 10 randomly selected structures with a $K^+$ ion passing the collar. The accessible molecular surface demonstrating occlusion or opening up of the conduction pathway at the Leu124 collar. The left and middle panels in the top row are reference slabs from the crystal structure, slightly enlarged over Fig. 2a. Side chains and surface of Leu124 are coloured yellow, lipidic alkyl chains orange, and potassium depicted as purple spheres. (**b**) Hexagonal bin plot of diagonal distances, D1 vs D2 (see Fig 2d for definition) for structures in which an ion is passing through the Leu124 collar extracted from unrestrained MD. Symmetry equivalent pairs are included in the count and representation. Structures were distributed into hexagonal bins on the basis of D1-D2 couplets. The count in each bin is described by a colour key, with dark orange representing the greatest number of structures and white the least. The predominant central peak is as in Fig 2d and reflects a mean pore diameter of approximately 9 Å. Using data from short simulations, each corresponding to ~490 ps either side of the Leu-Cδ 'plane', the couplets of structures at the precise moment of passing the plane are plotted as cyan dots. The symmetry-equivalent D2-D1 couplets are shown as black dots. This illustrates the range of aperture size $K^+$ is able to permeate.

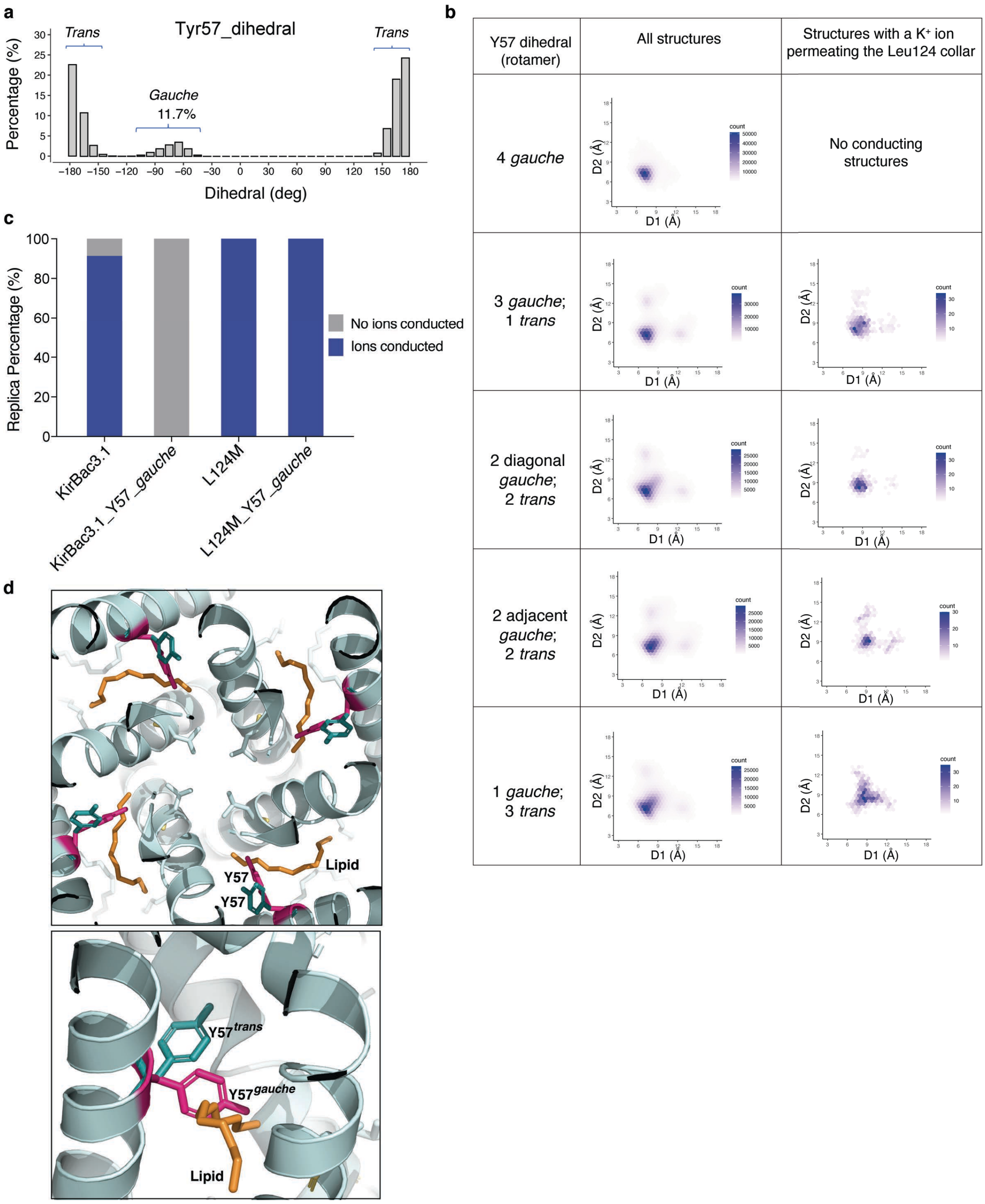


**Supplementary Figure 6. The *gauche* rotamer of Tyr57 blocks lipid entering the fenestrations.** (**a**) The proportion of the *gauche* conformer in the natural population, estimated by extracting data from all simulation structures, is close to 12 %. (**b**) Hexbin plots (as in Fig. 2d) showing the effect of excluding lipid from fenestrations on pore aperture. Notably, when all four Tyr57 adopt a *gauche* rotamer the collar remains small and impassable. (**c** Quantification of ion conduction of simulation replicates of wild type KirBac3.1 and the L124M mutant when fenestrations are either untouched or blocked (Y57 *gauche*). (**d**) The *trans* rotamer of Tyr57 (deep cyan) allows lipid (orange) to enter the fenestrations, whereas the *gauche* rotamer (magenta) blocks the fenestration and lipid. (Top) Viewed along the molecular 4-fold axis (*z*), perpendicular to the membrane. (Bottom) Viewed from within the plane of the membrane.

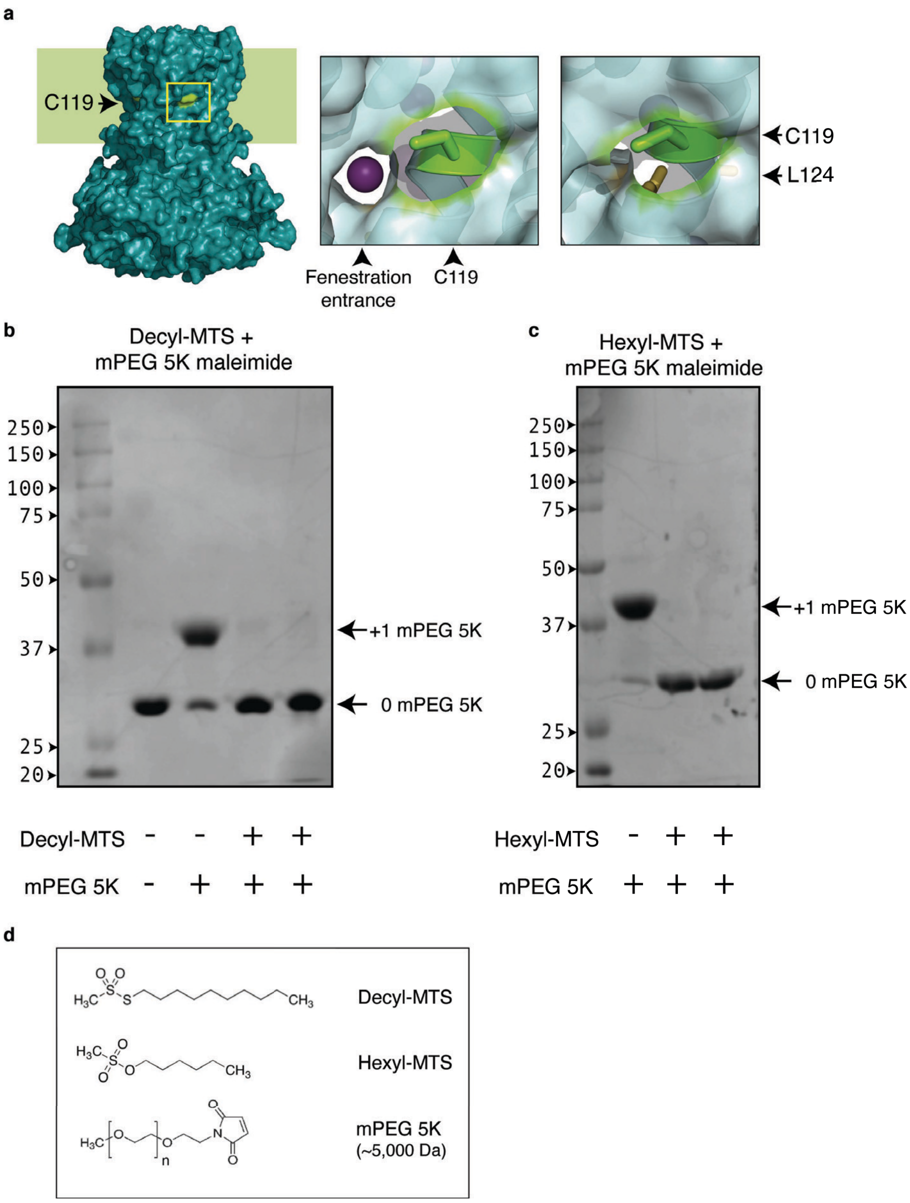


**Supplementary Figure 7. MTS-alkyl derivatization.** (**a**) The position of Cys119 (C119; chartreuse; in a yellow box) relative to fenestrations near the midpoint of the membrane is depicted by a molecular surface representation of KirBac3.1-SCS; the membrane region is shaded. Middle panel shows the spatial relationship of C119 to the fenestration entrance, aligned to show an internal $K^+$ ion through the fenestration; Right-hand panel shows the proximity to Leu124 The molecular surface immediately covering Cys119 is omitted for clarity. (**b**) **Gel-shift assay used to indicate completeness of** formation of C119-decyl by MTS-derivatisation of KirBac3.1-SCS, comparing the relative migration of unreacted (denatured; monomeric) KirBac3.1 with that of KirBac3.1 reacted with 5 mM methoxy PEG maleimide 5K and KirBac3.1 derivatized with decyl-MTS prior to exposure to 5 mM methoxy PEG maleimide 5K. The shift in the band migration in lane 3 corresponds to the addition of a single 5K PEG per subunit of KirBac3.1. The absence of a band shift in lanes 4 and 5 indicates decyl-MTS has reacted with Cys119. (**c**) Gel-shift assay for hexyl-MTS derivatization of KirBac3.1-SCS, comparing the relative migration of KirBac reacted with 5 mM methoxy PEG maleimide 5K and KirBac derivatized with decyl-MTS before exposure to 5 mM methoxy PEG maleimide 5K. (**d**) Chemical formulae as indicated.

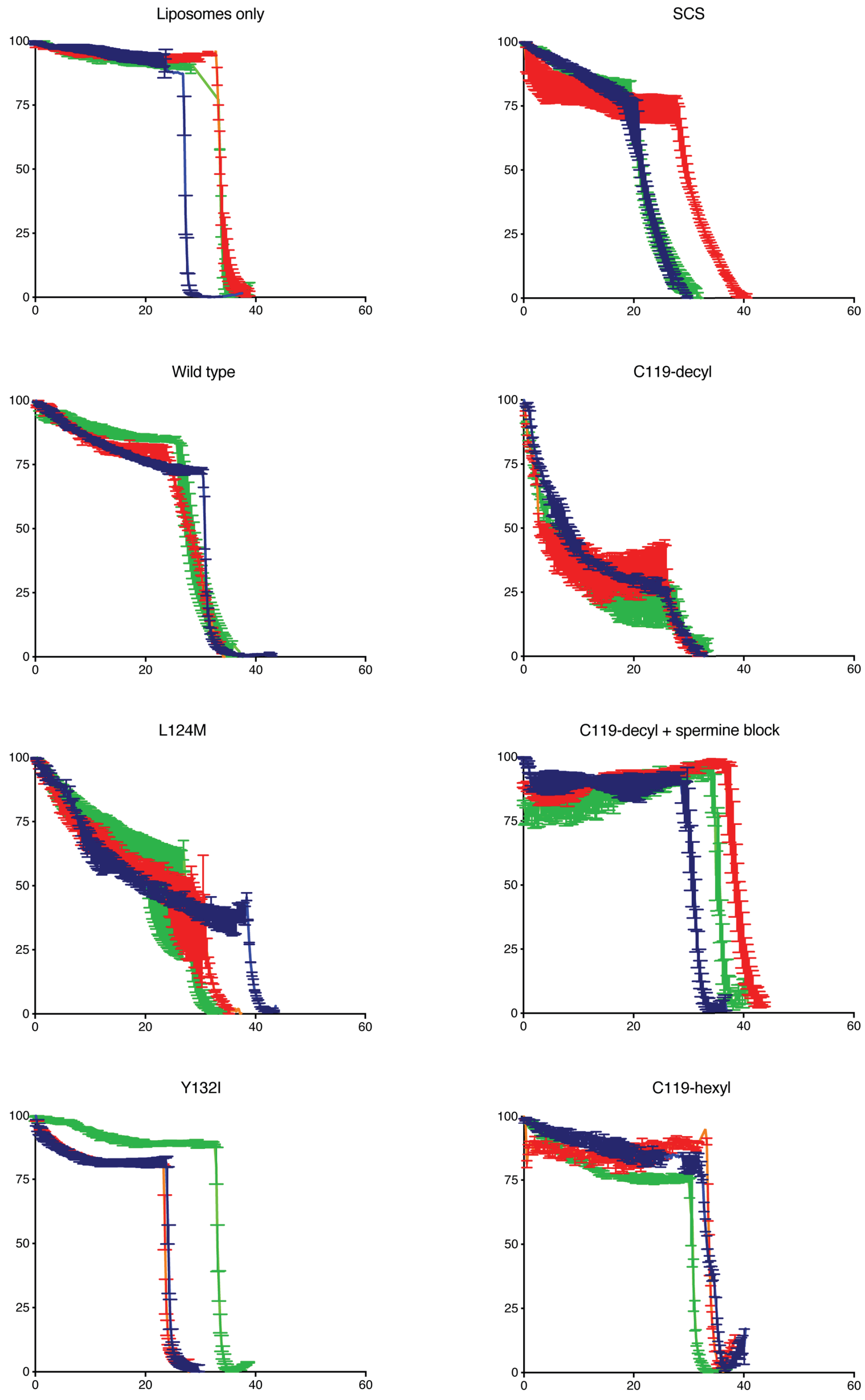


**Supplementary Figure 8. Normalised data from liposomal fluorescence flux (ACMA) assays**.
Three independent replicates are shown for each experimental condition, indicated by the title above each plot. For each experiment, each colour (red, blue, green) represents an independent reconstitution. The standard deviations (of three technical replicates) are indicated by error bars.

**a**

| | Po | Ps | Po/Ps | Pburst | Rate O -> S (ms$^{-1}$) | Rate S -> O (ms$^{-1}$) | Io (pA) | Is (pA) | Vm (mV) | Is/Io |
|---|---|---|---|---|---|---|---|---|---|---|
| C119-Decyl | | | | | | | | | | |
| mean, n=5 | 0.55 | 0.24 | 2.92 | 0.69 | 0.41 | 0.76 | -3.18 | -1.25 | -80 | 0.393 |
| sem | 0.08 | 0.05 | 1.07 | 0.06 | 0.15 | 0.15 | 0.35 | 0.23 | | 0.029 |
| C119-Octyl | | | | | | | | | | |
| n=1 | 0.36 | 0.41 | 0.89 | 0.46 | 1.25 | 1.07 | -3.56 | -0.76 | -86 | 0.213 |
| C119-Hexyl | | | | | | | | | | |
| mean, n=2 | 0.25 | 0.36 | 0.71 | 0.41 | 0.84 | 0.64 | -3.20 | -1.3 | -80 | 0.510 |
| range/2 | 0.11 | 0.17 | 0.03 | 0.01 | 0.05 | 0.05 | | | | 0.139 |

**b**

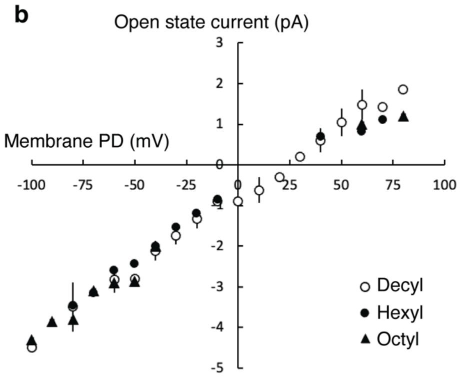


**c**

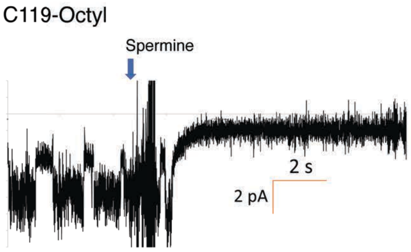


**Supplementary Figure 9. HMM analysis of single channel recordings of C119-alkylated KirBac3.1.** (**a**) Kinetic analysis, where I = current (pA), V = voltage, Po = probability of being fully open (maximal conductance), Ps = probability of being in a substate (lower conductance), Pburst is Po/(Po+Ps), the probability of being in the fully open state within a burst of activity, k = rate constant between fully open and substates. (**b**) Current-voltage relationship shows that aside from the differences in kinetics, the three derivatised species are fundamentally similar. (**C**) Spermine sensitivity of C119-octyl at -80 mV.

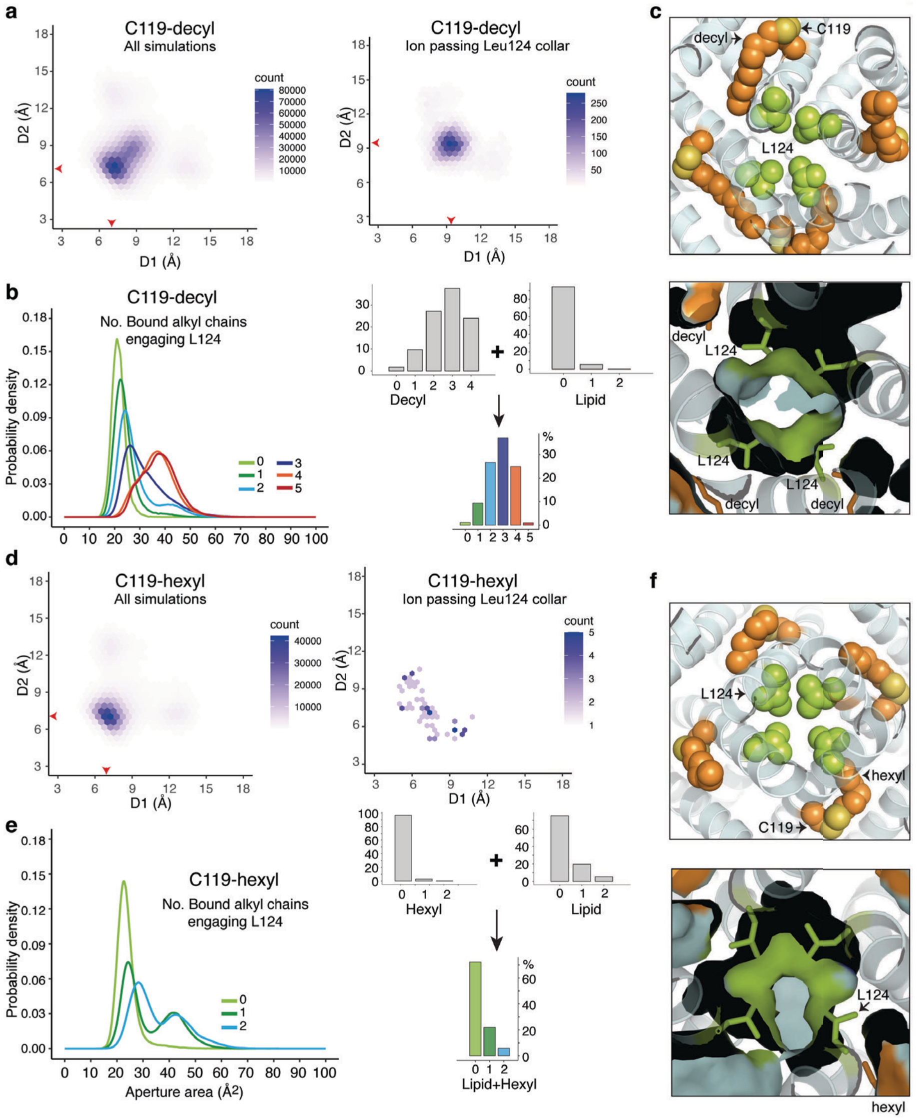


**Supplementary Figure 10. Aliphatic chain occupancy affects pore aperture in the C119-derivatised channels** similarly to the effect of lipids on wild type. C119-decyl and C119-hexyl substituents engaging Leu124, and the effect on pore aperture. (**a**) Hexagonal bin plots analysing the aperture diagonals in structures extracted from 70 simulations of unrestrained MD of C119-decyl. Distances are between centres of the red dots (midway between Cδ1 and Cδ2) in the schematic in Fig 2d. Red arrows on the plots indicate the population of highest density. (Left) Analysis of all structures. (Right) Analysis of structures in which a $K^+$ is passing the collar (**b**) The number of alkyl chains engaging with Leu124 side chains is correlated to the cross-sectional area of the 'open collar' between leucine side chains. (Right) In the C119-decyl channel, total alkyl occupancy can be contributed to by membrane lipids pushing in. (**c**) Transverse slice through the C119-decyl mutant at the Leu124 collar (yellow-green) of a random simulation structure with three fenestrations in which the occupying decyl substituents (orange) are engaging Leu124 residues. The accessible molecular surface below demonstrates the effect on the steric leucine cluster. (**d**) As for panel A*,* but hexbin plots are for C119-hexyl. Twenty simulations were carried out. (**e**) As for panel *b* but showing C119-hexyl distribution (**f**) Transverse slice through the C119-hexyl mutant as for panel C, shows one of four occupying hexyl substituents is reaching in far enough to contact a Leu124 residue (lower right corner)- the probability of this is only ~3 %. The accessible molecular surface below shows the sole leucine-hexyl contact allows limited opening of the steric cluster.

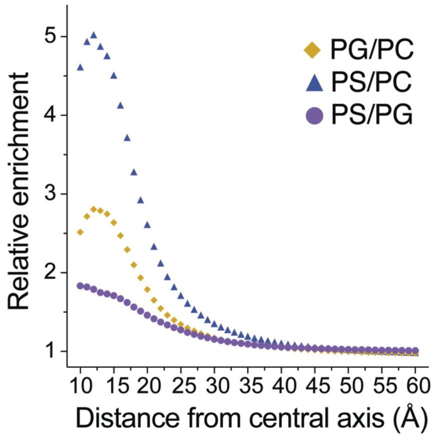


**Supplementary Figure 11.**

Pairwise cross comparison of the probability density distribution of PC, PS and PG lipid head groups surrounding the pore derived from the coarse grained simulation data, showing that (on average) both anionic lipids are more closely associated with the channel than PC.

**Supplementary Table 1.** Crystallographic Structure Determination and Refinement.

| **Data collection** | SCS<br>KirBac3.1_C71S-C119-C262S | L124M<br>KirBac3.1_C71S-C119-C262S_L124M |
|---|---|---|
| Processing system | XDS | XDS |
| Space group | $P42_12$ | $P42_12$ |
| Monomers per asymmetric unit | 1 | 1 |
| Cell dimensions (Å) | *a* 106.884, *c* 89.760 | *a* 106.577, *c* 89.597 |
| Wavelength (Å) | 0.9537 | 0.9537 |
| Resolution range (Å) | 47.80/2.40 | 47.663/2.708 |
| Observed/Unique reflections | 276,325/20,993 | 190,560/26,392 (Friedel pairs) |
| $R_{merge}$ (overall/outer shell) [a] | 0.235/2.425 | 0.158/1.305 |
| I/σI (overall/outer shell) | 8.60/0.52 | 10.52/1.35 |
| Completeness (overall/outer shell) | 99.8 /99.3 | 99.8/99.2 |
| Redundancy (overall/outer shell) | 13.16/11.84 | 7.22/6.86 |
| $CC_{1/2}$ (overall/outer shell)[b] | 99.8/40.4 | 99.8/57.9 |

| **Data refinement** | | |
|---|---|---|
| Program | PHENIX | PHENIX |
| Resolution range (Å) | 47.80 – 2.40 | 42.08-2.723 |
| Reflections (refine/test) | 20,950/1,050 | 14,342/700 |
| *GR* factor ($R_{cryst}/R_{free}$) | 0.2281/0.2583 | 0.2197/0.2507 |
| Wilson B-factor | 50.2 | 61.2 |
| Mean isotropic *B*-factor $B_{iso}$ (Å$^2$) | 55.7 | 60.1 |
| RMSD Bond lengths (Å) | 0.002 | 0.002 |
| RMSD Bond angles (°) | 0.51 | 0.48 |
| Ramachandran outliers | 0 | 0 |
| Rotamer outliers | 3 (1%) | 3 (1%) |
| PDB code | 7N9L | 7N9K |

[a] $R_{merge} = \sum | I_{obs} - I_{calc} | / \sum I_{obs}$

[b] $CC_{1/2}$ = percentage of correlation between intensities from random half-datasets. Outer shell values are significant at the $p$=0.1% level[1].

**Supplementary Table 2.** Summary of Molecular Dynamics Simulations.

| **Equilibration** | | |
|---|---|---|
| Channel condition | | Simulation length (ns) |
| Wild Type | | 101.5 |
| Y132I | | 101.5 |
| L124M | | 101.5 |
| L124M: 4 *gauche* | | 101.5 |
| Y57: 4 *gauche* | | 101.5 |
| Y57: 3 *gauche*; 1 *trans* | | 101.5 |
| Y57: 2 diagonal *gauche*; 2 *trans* | | 101.5 |
| Y57: 2 adjacent *gauche*; 2 *trans* | | 101.5 |
| Y57: 1 *gauche*; 3 *trans* | | 101.5 |
| C119-decyl | | 101.5 |
| C119-hexyl | | 101.5 |
| **Steered Molecular Dynamics Pulling** | | |
| Channel condition | Pulling Speed (Å/ns) | Simulation length (ns) |
| Wild Type | 0.2 | 150 |
| Y132I | 0.2 | 120 |
| L124M | 0.2 | 110 |
| Y57: 4 *gauche* | 0.2 | 110 |
| **Umbrella Sampling** | | |
| Channel condition | Windows number * Simulation length per window (ns) | Simulation length (ns) |
| Wild Type | 74*310 | 22,940 |
| Y132I | 97*200 | 19,400 |
| L124M | 53*310 | 16,430 |
| Y57: 4 *gauche* | 52*310 | 16,120 |
| **Unrestrained MD** | | |
| Channel condition | Simulation number * Simulation length (ns) | Simulation length (ns) |

| | | |
|---|---|---|
| Wild Type | 70*200 | 14,000 |
| C119-decyl | 70*200 | 14,000 |
| C119-hexyl | 20*200 | 4,000 |
| Y57: 4 *gauche* | 20*200 | 4,000 |
| Y57: 3 *gauche*; 1 *trans* | 20*200 | 4,000 |
| Y57: 2 diagonal *gauche*; 2 *trans* | 20*200 | 4,000 |
| Y57: 2 adjacent *gauche*; 2 *trans* | 20*200 | 4,000 |
| Y57: 1 *gauche*; 3 *trans* | 20*200 | 4,000 |
| L124M: 4 *gauche* | 20*200 | 4,000 |
| **Total simulation length (ns)** | | **151,718** |
| **Coarse Grained Simulations** | | |
| **Channel Condition** | **Simulation length (ns)** | **Simulation number * Simulation length (ns)** |
| Wild Type | 5*20000 | 100,000 |
| **Total coarse grained simulation length (ns)** | | **100,000** |

**Supplementary Table 3.** Summary statistical data on fluorescence change for functional **ACMA** assays for control, wildtype, mutant, and C119-alkyl KirBac3.1.

| Condition | Mean fluorescence change prior to valinomycin addition ± SEM | Independent replicates (technical replicates for each experiment) | Significance compared to KirBac3.1 | | Significance compared to KirBac3.1-SCS | |
|---|---|---|---|---|---|---|
| Liposomes only (channel free) | 7.74 ± 0.81 | 10 (5, 3, 3, 5, 4, 4, 4, 4, 3, 3) | **** | <0.0001 | **** | <0.0001 |
| Wildtype KirBac3.1 | 27.99 ± 2.65 | 6 (5, 3, 4, 3, 3, 3) | | | ns | 0.6941 |
| KirBac3.1-SCS ( *i.e.* C71S, C262S) | 29.71 ± 3.34 | 3 (3, 3, 3) | ns | 0.6941 | | |
| L124M | 55.71 ± 6.30 | 5 (3, 3, 3, 3, 3) | ** | 0.0019 | * | 0.0449 |
| Y132I | 16.81 ± 1.72 | 3 (3, 3, 3) | * | 0.0289 | * | 0.0265 |
| C119-Decyl | 77.22 ± 2.36 | 3 (6, 3, 3) | **** | <0.0001 | **** | <0.0001 |
| C119-Hexyl | 16.86 ± 3.61 | 5 (3, 3, 3, 3, 3) | * | 0.0329 | ns | 0.0649 |
| C119-Decyl + Spermine block (500 μM / 1000 μM) | 5.56 ± 4.12 | 3 (3, 6, 6) | ** | 0.0021 | ** | 0.0037 |

**Supplementary Table 4.** Summary rates data for ACMA assays.

| Condition | Mean rate (time constant, tau) ± SEM (min) | Independent replicates (technical replicates for each experiment) | Significance compared to Wildtype | | Significance compared to SCS | |
|---|---|---|---|---|---|---|
| Wildtype KirBac3.1 | 20.4 ± 2.34 | 6 (5, 3, 4, 3, 3, 3) | | | ns | 0.8041 |
| KirBac3.1-SCS | 19.1 ± 3.23 | 3 (3, 3, 3) | ns | 0.8042 | | |
| L124M | 19.6 ± 2.89 | 5 (3, 3, 3, 3, 3) | ns | 0.8580 | ns | 0.9269 |
| Y132I | 11.9 ± 7.49 | 3 (3, 3, 3) | ns | 0.1198 | ns | 0.2480 |
| C119-Decyl | 5.22 ± 1.2 | 3 (6, 3, 3) | ** | 0.0087 | * | 0.0322 |
| C119-Hexyl | 15.10 ± 3.23 | 5 (3, 3, 3, 3, 3) | ns | 0.2786 | ns | 0.4867 |
